# IFNγ at the crossroads of systemic protection and brain vulnerability to coronavirus infection

**DOI:** 10.64898/2026.07.17.739144

**Authors:** Agustina Sabater, Ana P. Arevalo, Paula Perbolianachis, Pablo Sanchis, Gaston Pascual, Jorge L. Porfido, Rocio Seniuk, Madelón Portela, Maria P. Valacco, Javier Cotignola, Elba Vazquez, Estefania Labanca, Juan Manuel Carballeda, Pilar Moreno, Gonzalo Moratorio, Martina Crispo, Geraldine Gueron, Ayelen Toro

## Abstract

The COVID-19 pandemic, together with previous outbreaks caused by highly pathogenic coronaviruses, highlighted the importance of understanding the host determinants that influence pathogenesis, both to improve our understanding of disease mechanisms, and to strengthen preparedness for future outbreaks. In our previous work we demonstrated dysregulation of IFNγ response genes in COVID-19 patients, associated with both age and viral burden, positioning IFNγ as a key mediator of host defense against SARS-CoV-2. Here, we used a preclinical murine model of coronavirus infection based on murine hepatitis virus (MHV-A59), widely validated for studying SARS-CoV-2 pathogenesis, together with IFNγ knockout (IFNγ-KO) mice to investigate the role of this cytokine during infection. We performed a comprehensive morphological, biochemical, hematological and proteomic characterization of infected wild-type (WT) and IFNγ-deficient mice. Plasma proteomics revealed impaired inflammatory and coagulation-related responses in IFNγ-KO animals compared to WT controls. We found increased viral load and infectious particles in peripheral organs, including liver, spleen, heart and muscle in IFNγ-KO mice, whereas these were significantly reduced in the brain. Region-specific analysis further demonstrated decreased viral load in the prefrontal cortex and hippocampus of IFNγ-KO animals. Consistently, bioinformatics analysis of transcriptomics data of IFNγ-treated primary neuron cultures, together with frontal cortex from human COVID-19 patients, revealed activation of neuroinflammatory and neurological disease-associated pathways, supporting a role for IFNγ in driving brain inflammatory responses during coronavirus infection. Together, these findings reveal a dual role for IFNγ during coronavirus infection: it is important for controlling systemic viral dissemination and limiting peripheral tissue damage, yet it may also promote viral susceptibility and exacerbated inflammatory responses in the brain. These results support IFNγ modulation as a potential therapeutic strategy to prevent or mitigate neurological complications associated with COVID-19.

**Author Summary:** Viral infections are controlled by the immune system, but the same responses that protect the body can sometimes contribute to tissue damage and disease. We investigated how a key immune molecule, called Interferon Gamma, influences the outcome of Coronavirus infection in different parts of the body. Using a mouse model of Coronavirus infection with Interferon Gamma deficiency, we found that this molecule plays two opposing roles: it helps control the spread of the virus throughout the body, but it also increases vulnerability of the brain to infection and inflammation. When this immune signal was absent, animals showed higher amounts of virus in several peripheral organs but surprisingly had lower levels of virus in specific brain regions. We further found that this immune response was associated with changes in inflammation and neurological processes in human data. Our findings reveal that antiviral immunity is not always uniformly protective and that its effects depend on the tissue involved. Understanding this balance may help guide future approaches to reduce virus-associated complications while preserving the protective functions of the immune response.

## Introduction

Coronaviruses constitute a family of viruses accountable for various afflictions, including the common cold, severe acute respiratory syndrome (SARS), and Middle East respiratory syndrome (MERS). They infect humans, other mammals and avian species, entailing not only a challenge for public health but also a veterinary and economic concern (1).

The main therapeutic avenues to halt viral infection consist of targeting the virus directly or boosting the host immune system. Thus, deciphering the molecular and cellular mechanisms of coronavirus infection, as well as host determinants in the disease, is critical for the development of effective intervention strategies (1). In line with this, interferons (IFNs) provide a first line of host defense against viral infections by promoting an intracellular environment that restricts viral replication. In the local environment, IFNs promote the transcription of hundreds of interferon-stimulated genes (ISGs), encoding proteins with antiviral, antiproliferative and immunomodulatory activities, and work in synergy to inhibit viral replication via multiple mechanisms (2,3). The IFN family includes three main classes of related cytokines: type I IFNs (IFN-I), type II IFN (IFN-II) and type III IFN (IFN-III). There are many IFNs-I (IFNα, IFNβ, IFNδ, etc.) and many IFNs-III (different IFNλs); however, there is only one IFN-II: IFNγ. Particularly, IFNγ binds to the ubiquitously expressed IFNγ receptor (IFNGR), activating a downstream signaling pathway that subsequently regulates gene expression via the gamma interferon-activated site (GAS) of ISGs (4).

While limited evidence links IFN-II to COVID-19, serum levels of IFNγ were significantly elevated in symptomatic patients compared with asymptomatic COVID-19 groups (5). Additionally, in a murine preclinical model it was reported that infection with MHV-A59, a well-established betacoronavirus surrogate for SARS-CoV-2 (6), was associated with an increment of IFNγ in plasma (7). Our previous work highlighted the relevance of interferon-mediated responses in the context of coronavirus infection. Briefly, we described the upregulation of the interferon-stimulated gene *MX1* (8) in response to viral infection and its potential contribution to antiviral defense mechanisms. Furthermore, we identified a significant dysregulation of genes associated with the IFNγ signaling pathway in COVID-19 patients, demonstrating that their expression correlated with both viral load and patient age, thereby positioning IFNγ as a central component of the host antiviral response and severity (9).

Building on these findings, we hypothesize that IFNγ–driven programs are critical determinants of disease progression and host response. In this study, we functionally dissect the role of IFNγ during coronavirus infection by leveraging a preclinical IFNγ-knockout (KO) mice model infected with Murine Hepatitis Virus (MHV). Our work delves into the details of how IFNγ shapes tissue-specific viral dissemination and pathogenesis. We demonstrated that IFNγ-deficiency is associated with increased viral dissemination to peripheral organs but decreased in RBCs and brain, particularly in the prefrontal cortex and hippocampus, revealing a previously unrecognized role for IFNγ in neurotropism. Moreover, by transcriptomics and proteomics analysis, we identified neuroinflammatory and neurological-disease-associated programs that are dysregulated in the absence of IFNγ, upon IFNγ treatment of neuron cultures, and in the frontal cortexes of COVID-19 patients. These results point out to a robust association between IFNγ and coronavirus-induced neurological pathology.

## Methods

### Virus and cells

Murine Hepatitis Virus (MHV-A59, ATCC VR-764) was cultured in laboratory conditions by performing 10 serial passages in murine L929 cells (SIGMA). This stock was stored at −80 °C until further use. Cells were maintained in DMEM medium (Gibco) supplemented with 10% vol/vol FBS (Fetal Bovine Serum, Gibco), 1% vol/vol penicillin-streptomycin and incubated at 37 °C and 5% CO_2_.

### Animals

C57BL/6J and the transgenic strain B6.129S7-*Ifng^tm1Ts^*/J (10) (#000664 and #002287, The Jackson Laboratory, ME, USA) were bred at the animal facility of the Laboratory Animals Biotechnology Unit of Institut Pasteur de Montevideo under the conditions described in (9). The mutant genotype was periodically checked by PCR genotypification. All the experimental protocols were approved by the institutional Comisión de Ética en el Uso de Animales (protocol #006-22) and were performed according to national law #18.611 and relevant international laboratory animal welfare guidelines and regulations.

### *In vivo* experiments

C57BL/6J WT and IFNγ-KO female mice (8–10 weeks old) were weighted and randomly distributed into four groups according to the experiment: (a) mock-infected wild-type (WT, n = 9), (b) MHV-infected wild-type (MHV WT, 6,000 PFU *i.p.*, n = 15), (c) mock-infected IFNγ-KO (IFNγ-KO, n = 5), or d) MHV-infected IFNγ-KO (MHV IFNγ-KO, 6,000 PFU *i.p.*, n = 20). Mock-infected animals were injected *i.p.* with 100 µL of vehicle (PBS). Five days post-infection, mice were weighted, bleed and euthanized by deep anesthesia and cervical dislocation. Blood and tissue samples were collected as previously described in (7). Sample size was estimated using power analysis considering the expected effect size, variability observed in preliminary data, an alpha parameter of 5%, and a statistical power of 80% (7).

For baseline characterization, blood samples were collected from all animals used in this study before the infection protocol, and organs were harvested from mock-infected controls after the experiments, in order to minimize the number of animals used, aligned with the 3Rs principle.

### Dissection of brain regions

For the analysis of different brain regions, 5 C57BL/6J WT and 5 IFNγ-KO female mice (8–10 weeks old) were infected with MHV-A59 (6,000 PFU) *i.p.* Five days after the infection, mice were euthanized by deep anesthesia and cervical dislocation, and the brains were quickly removed. Hippocampus, prefrontal cortex, cerebellum and hypothalamus were dissected using a razor blade. The dissected tissues were flash frozen in liquid nitrogen for RNA extraction.

### Organ processing

Organs were weighted upon necropsy. Then, samples from liver, lung, brain, heart, kidney, spleen and quadricep muscle were retrieved and excess blood was removed by washing twice with PBS. A piece of each organ was frozen in TRIzol (Invitrogen) in liquid nitrogen for RNA extraction and another was further processed for functional viral particles assessment.

### Blood processing

#### Biochemical parameters

Individual whole blood (100 μL) was collected in 20 U/mL of heparin (Heparin Sodium salt, SIGMA) and analyzed for liver and kidney biochemical profile using the Pointcare V2 automatic device (Tianjin MNCHIP Technologies Co, China) at the beginning (pre-infection determination) and at the end of the experiment (post-infection determination). Analyzed parameters included total protein (TP), albumin (ALB), globulin (GLO), aspartate aminotransferase (AST), alanine aminotransferase (ALT), gamma-glutamyl transpeptidase (GGT), blood urea nitrogen (BUN), and creatinine (CRE). ALT detection limit corresponds to 1500 (U/L) and AST to 1600 (U/L).

#### Hematological parameters

For hematological analysis, 20 μL of blood were collected into 0.5 mL microtubes containing EDTA potassium salts (W anticoagulant, Wiener lab, Santa Fe, Argentina) in a ratio of 1:10 (EDTA: blood) before and after infection. All measurements were conducted within four hours after collection. Red blood cells (RBC) count, hemoglobin (HGB), hematocrit (HCT), white blood cells (WBC), platelets (PLT) counts, and immune populations (lymphocytes, neutrophils, basophils, monocytes and eosinophils) percentages were evaluated using the auto hematology analyzer BC-5000Vet (Mindray Medical International Ltd, China). Neutrophil-to-lymphocyte ratio (NLR) and platelet-to-lymphocyte ratio (PLR) were calculated using absolute values.

#### Viral and proteomics analysis

Another aliquot of individual whole blood (100 μL) was collected in 10% vol/vol EDTA and was centrifuged at 3,000 × g for 30 min at 4 °C to separate RBC and plasma fractions. 100 μL of PBS 1x were added to each fraction and split into three aliquots. One aliquot of each fraction was used to perform TRIzol RNA extraction for viral genome equivalents determination, another aliquot was used to determine viable viral particles by plaque assay, and a third one was subjected to proteomics assays.

### Viral genome equivalents in organs and blood fractions

Total RNA was isolated using TRIzol reagent according to the manufacturer’s protocol. The extracted RNA was used for MHV-A59 detection using a specific sensitive qPCR protocol using TaqMan Fast Virus 1-Step Master Mix (Applied Biosystems) with a Taq-man probe (ACAAGCTCAGGCACCTCCTGTACAA) labeled at the 5′-end with FAM (10 mM). The following primers were used for viral load determination: *Nsp2* forward: 5’-TGGATGGCTTTGCTACCAG-3’, and *Nsp2* reverse: 5’-CCAGACAAGATAGAAACCGAC-3’. All RT-qPCRs were performed in duplicate. Thermal cycling was run on a QuantStudio™ seven (Applied Biosystems). For the viral load estimation, we used a standard curve as previously described (11), with the equation being y = -3.7x + 46.1 and the coefficient of determination (R^2^) equal to 0.9917.

### Median tissue culture infectious doses

Infectious particles were assessed in supernatants obtained from different MHV-infected mice organ samples by the Median Tissue Culture Infectious Doses (TCID_50_) technique. A total of 5 × 10^4^ L929 cells (SIGMA) were seeded in 96-well plates. Tenfold serial dilutions of the supernatants were prepared in serum-free DMEM media (Gibco). Infections were performed in twelve replicates. After five days, living cell monolayers were fixed and stained with crystal violet 0.2% vol/vol in formaldehyde 10% vol/vol. TCID_50_ values obtained were normalized to the weight of the organ sampled and expressed as TCID_50_/g of tissue.

### Plaque assays

Infectivity was assessed in blood fractions using plaque assays. L929 cells were seeded in 12-well plates and supernatants from MHV-infected mice blood fractions (plasma or RBC-enriched) were serially ten-fold diluted in DMEM serum-free medium (Gibco). After 1 h of viral adsorption, monolayers were topped with a semisolid overlay of DMEM medium supplemented with 2% FBS and 1% WT/vol agarose. Forty-eight hours post infection, cells were fixed with formaldehyde 10% vol/vol and stained with crystal violet 0.2% vol/vol.

### Proteomics analyses

For proteomics analysis, 5 C57BL/6J WT and 5 IFNγ-KO female mice (8–10 weeks old) were infected with MHV-A59 (6,000 PFU) *i.p*. Five days after the infection, blood samples were collected and blood fractionation was performed as mentioned in the *Blood processing* section. Plasma fractions were depleted from albumin, transferrin and G immunoglobulins using Multiple Affinity Removal Spin Cartridge Mouse-3 (Agilent Technologies).

Protein samples allowed to enter up to 1 cm into an SDS-PAGE resolving gel, fixed and stained with Coomassie G-250. In-gel protein digestion and peptide extraction were performed as previously described in (11). Tryptic peptides were analyzed on a nano-HPLC UltiMate 3000 (Thermo Scientific) coupled to an Orbitrap Exploris 240 mass spectrometer (Thermo Scientific) through an Easy-Spray source (Thermo Scientific) (11).

PatternLab for Proteomics software (http://www.patternlabforproteomics.org) (12,13) was used for peptide identification and label-free quantitative analysis. MS raw files were searched against a target-reverse database containing sequences from *Mus musculus* downloaded from UniProt (http://www.uniprot.org, 17/10/2023) and a database containing the most common contaminants in proteomics experiments. For data search, m/z precursor tolerance was set at 35 ppm. Methionine oxidation and cysteine carbamidomethylation were defined as variable and fixed modifications respectively. A maximum of 2 missed cleavages and 2 variable modifications per peptide were allowed. Search results were filtered by the Search Engine Processor (SEPro) algorithm from PatternLab with a maximum FDR value ≤ 1% at protein level and 10 ppm tolerance for precursor ions.

### Bioinformatics analysis

#### Datasets

We performed differential expression analysis using the data from GSE188847 (14,15) at Gene Expression Omnibus repository. This dataset contains RNA-seq profiling data from the frontal cortex of 22 COVID-19 patients and 23 uninfected controls. It also contains transcriptomic data from three independent primary cultures of human neurons treated with IFNγ 1 µg/mL (n=3) or vehicle (n = 3) for 72 h.

#### Differential Expression Analysis

Differential gene expression was performed using the Limma package (Linear Models for Microarray Analysis, version 3.58.1) (16) in R to study differential expressions both from proteomics data and RNA-seq data. In the case of non-normalized data, quantile normalization was applied (17). For proteomics data, only the genes observed in common for both genotypes were included for analysis. For RNA-seq data, the voom function in the Limma package was used for processing (18). For each available gene/protein, the fold changes (FCs) between conditions were calculated and expressed as log2FC. Benjamini-Hochberg correction was used to calculate the False Discovery Rate (adjusted P value).

#### Ingenuity Pathway Analysis

Ingenuity Pathway Analysis (19) (QIAGEN IPA, QIAGEN Inc., https://digitalinsights.qiagen.com/IPA) was used to summarize the differences and to study canonical and disease pathways with similar alterations. This software utilizes a comprehensive knowledge base of curated biological interactions and functional annotations, and maps the data into known biological networks, enabling the identification of regulators, pathways, and biological processes relevant to the experimental data. This approach facilitates a deeper understanding of the biological mechanisms underlying the observed molecular changes. We uploaded the differential expression analysis results comparing the quantitative proteomics data from plasma and RBC fractions of MHV IFNγ-KO *vs.* MHV WT mice, the differentially expressed genes on COVID-19 *vs.* control frontal samples and IFNγ-treated neurons *vs.* control. IPA calculates the p-value of overlap using the right-tailed Fisher’s Exact Test to identify significant pathways.

### Gene Expression Assays

RNA samples were depleted from DNA using DNase I (Thermo Scientific) and cDNAs were synthesized with LunaScript RT SuperMix Kit (New England Biolabs). FastStart Universal SYBR Green Master (Rox) (Roche) was used for real-time PCR amplification in a QuantStudio 3 Real-Time PCR System (Thermo Scientific), using the primers listed in **Table 1**. *Gapdh* was used as the internal reference gene. Data were analyzed using the method of 2^−ΔΔCT^ (20).

**Table 1.**
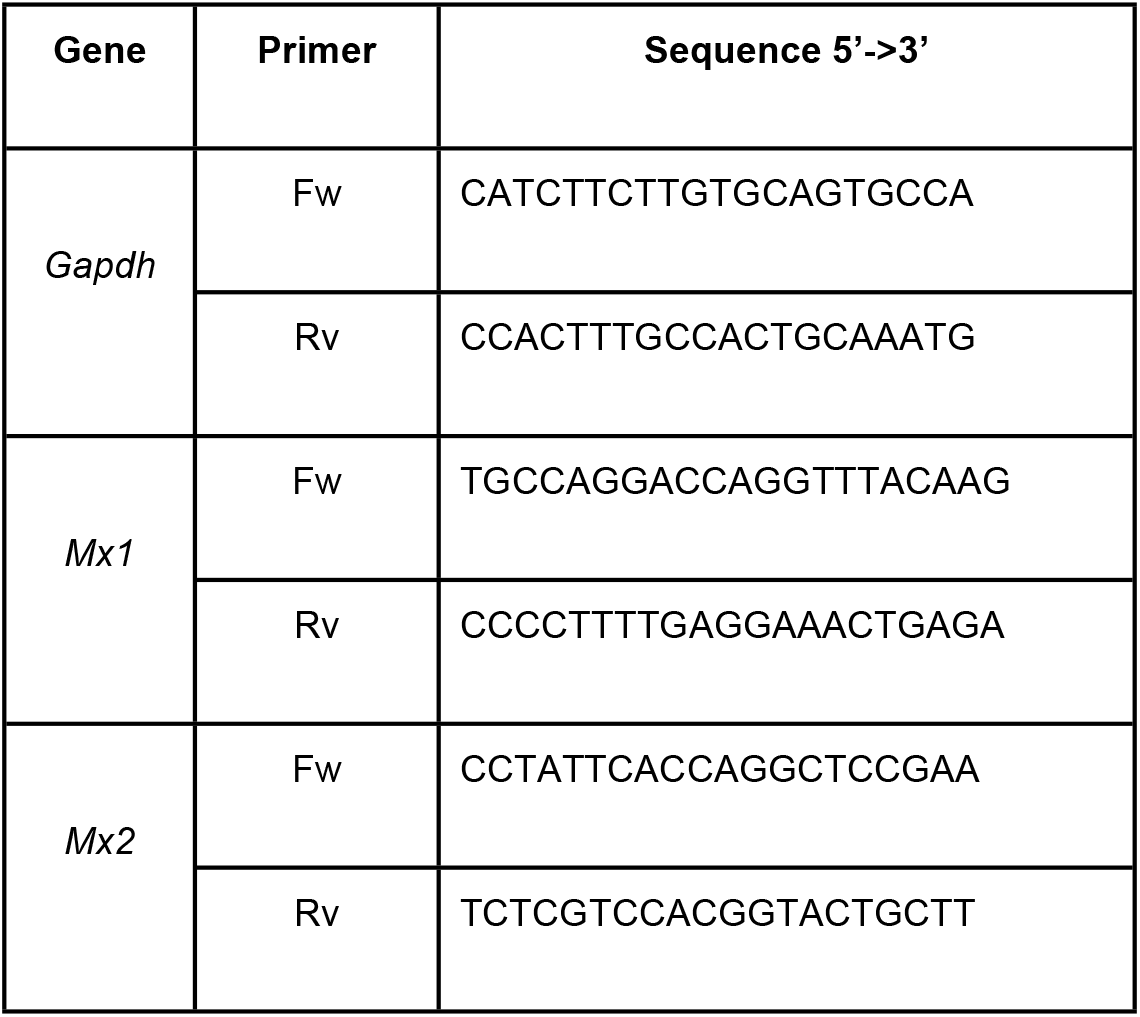
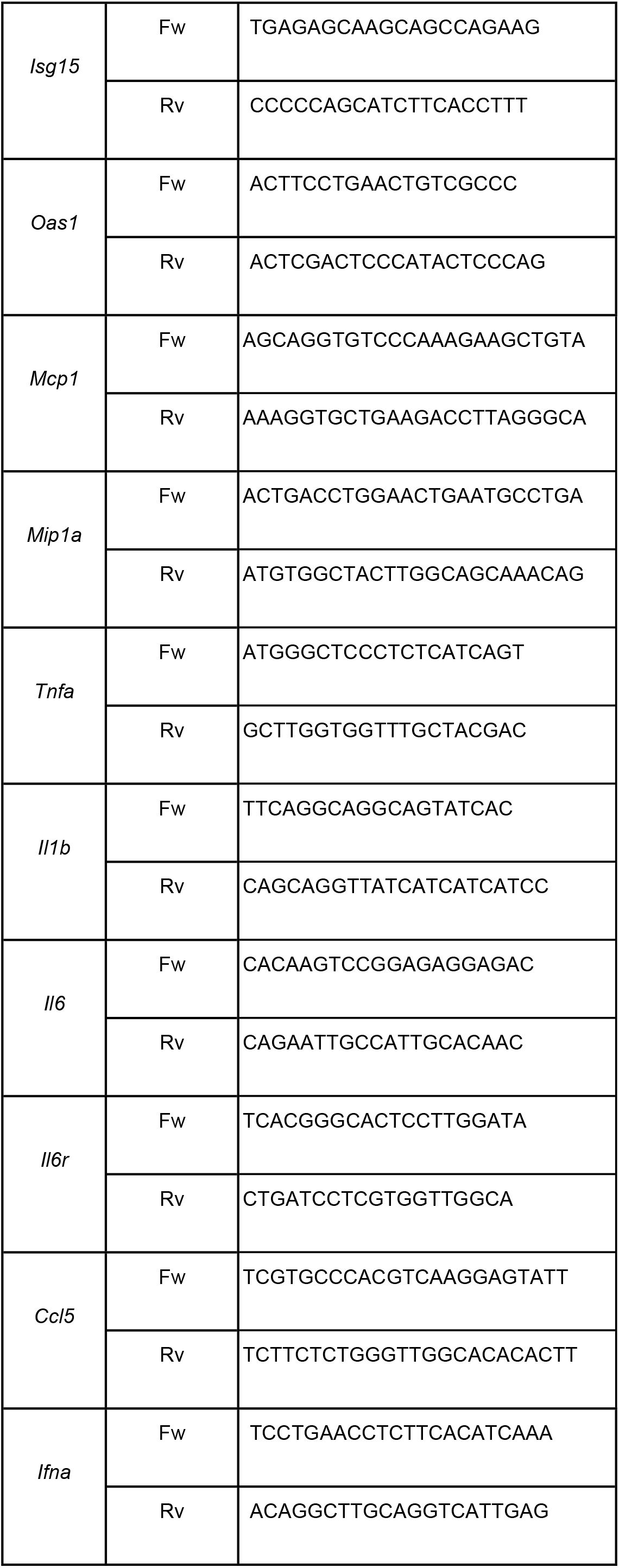
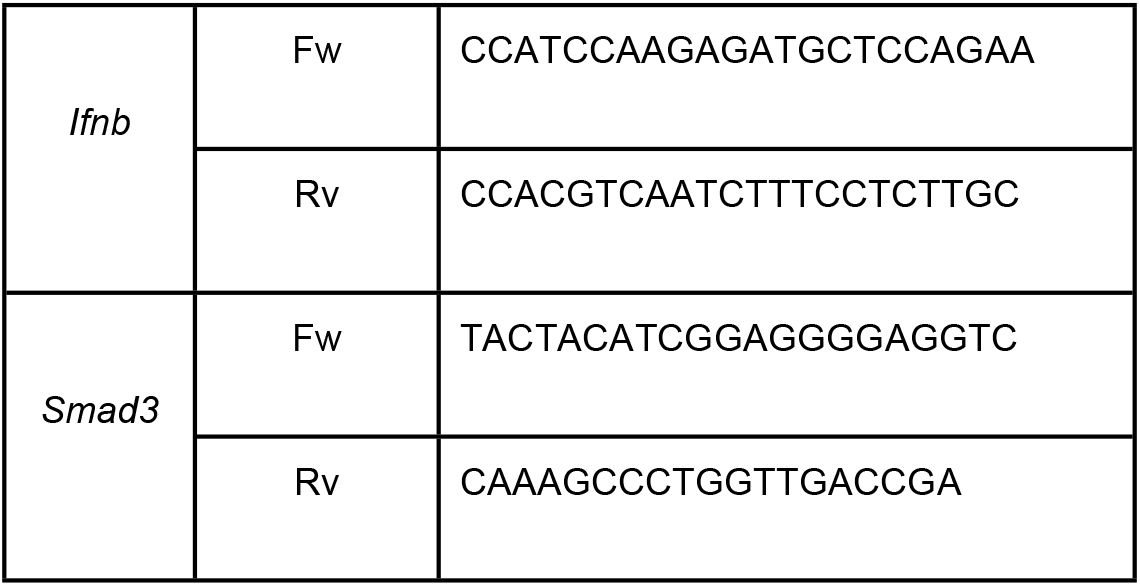
Primer sequences.

### Statistical analysis

Paired and unpaired student’s t-test was performed to determine statistical differences between pre- and post-infection measurements in mice or between genotypes. Post/pre ratios were used for assessment of changes in the magnitude of the infection-induced response, and unpaired student’s t-test was performed. One-way ANOVA followed by Tukey’s post-hoc test was used when comparing more than 2 groups. Mixed-effects analysis followed by Fisher’s Least Significant Difference (LSD) was performed to determine statistical differences between groups with pre and post conditions. Plots were generated and statistics were calculated using GraphPad Prism 11 software. Statistical significance was set at p < 0.05.

## Results

### Baseline characterization of the IFNγ-KO model

Based on our previous study where we identified IFNγ as a key host factor associated with coronavirus severity (9), we investigated the biological consequences of IFNγ deficiency using an IFNγ-KO mouse model. We first performed a comprehensive baseline morphometric, biochemical and hematological characterization of the IFNγ-KO strain to assess the intrinsic phenotype before the infection and compare it to WT mice (**Supplementary Fig. S1A**). No significant differences were observed in overall body weight between uninfected WT and IFNγ-KO animals (**Supplementary Fig. S1B**). Organ weight analysis revealed differences in liver weight, which was lower in IFNγ-KO mice (p < 0.05), while no differences were observed for spleen, lung, kidney, heart or brain (**Supplementary Fig. S1C**).

Biochemical assessment showed comparable baseline hepatic parameters, including total proteins, albumin, globulin, AST, ALT and GGT (**Supplementary Fig. S1Di**), and renal function markers such as blood urea nitrogen (BUN) and creatinine (CRE), between genotypes (**Supplementary Fig. S1Dii**). Regarding baseline immune cell quantification, although WBC total count did not differ across genotypes (**Supplementary Fig. S1Ei**), IFNγ-KO mice presented increased lymphocytes (p < 0.05), basophils (p < 0.01) and eosinophils (p < 0.01) proportion, while neutrophils were decreased (p < 0.05) (**Supplementary Fig. S1Eii**).

Hematological profiling demonstrated no specific changes between genotypes in baseline RBC count, hematocrit or hemoglobin levels (**Supplementary Fig. S1F**). However, IFNγ-KO mice presented increased platelet number (p < 0.05) and a decreased neutrophils-to-lymphocytes ratio (NLR) (p < 0.05) (**Supplementary Fig. S1F**).

Overall, the main differences between genotypes include different proportions of immune cells and platelets count. This starting point characterization provides the necessary reference framework to accurately interpret the subsequent infection-related outcomes in the IFNγ-KO model. Of note, this is the first in depth characterization performed for this strain.

### IFNγ deficiency affects systemic disease upon MHV infection

We next evaluated the impact of IFNγ deficiency in the context of MHV infection, aiming to define the contribution of IFNγ to the antiviral response against coronaviruses. For this aim, the characterized WT and IFNγ-KO animals were infected *i.p.* with MHV-A59 (6,000 PFU) and, after 5 days, mice were humanely euthanized and necropsy was performed (**Fig. 1A** and Supplementary Fig. S2A).

**Fig. 1.**
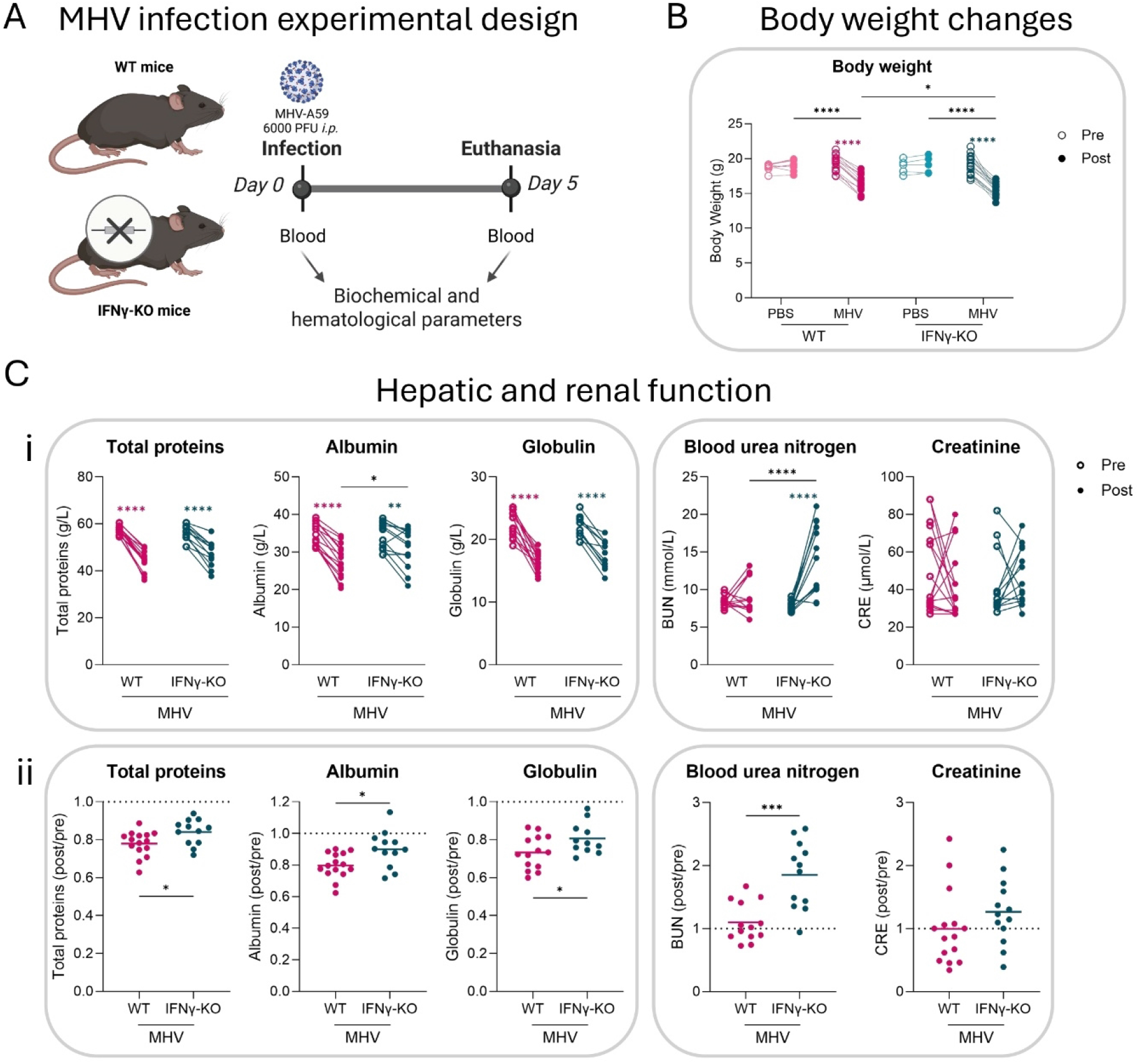
Systemic effects of MHV infection in WT and IFNγ-KO mice. **(A)** Schematic representation of the experimental design. Briefly, C57BL/6J wild-type (WT) and IFNγ-KO mice were infected i.p. with murine hepatitis virus (MHV) or mock (PBS). After 5 days, mice were humanely euthanized and necropsy was performed. Blood was collected at baseline (pre) and post-infection (post). Biochemical and hematological parameters were analyzed. Created with BioRender.com. **(B)** Dot plot depicting body weight (g) pre and post MHV or mock infection of WT and IFNγ-KO MHV-infected and mock-infected mice. **(C)** Dot plots depicting the biochemical assessment of hepatic (total proteins (g/L), albumin (g/L) and globulin (g/L)) and renal (blood urea nitrogen (mmol/L) and creatinine (µmol/L)) function pre and post infection **(i)** and post/pre ratios **(ii)** in WT and IFNγ-KO MHV-infected mice. A post/pre ratio of 1 indicates no changes with the infection; ratios higher than 1 represent increases after infection; ratios lower than 1 represent decreases after infection. Comparing ratios between groups allows assessment of changes in the magnitude of the infection-induced response. Statistical significance was determined by mixed-effects analysis followed by Tukey’s **(Bi)** or Fisher’s LSD **(Ci)**, or by Student’s t test **(ii)**, and was set at p < 0.05, with *p < 0.05; **p < 0.01; ***p < 0.001; ****p < 0.0001.

Although no differences were observed in the baseline weight of WT and IFNγ-KO mice, MHV infection generated weight loss in both WT and IFNγ-KO mice (p < 0.0001, **Fig. 1B**). However, body loss was greater in IFNγ-KO mice compared to WT controls (16.4% loss for WT vs. 18.97% loss for IFNγ-KO, p < 0.05) (**Fig. 1B**). Regarding organ weight changes, MHV infection increased spleen weight in the WT group (p < 0.05) and lung weight in IFNγ-KO animals (p < 0.001), whereas kidney and brain presented higher weights in both cases (p < 0.0001 and p < 0.01, respectively), without differences between genotypes (**Supplementary Fig. S2B**). No significant differences were observed neither in heart weight among groups nor in liver weight between infected groups (**Supplementary Fig. S2B**). Hematological analysis did not show differences in RBC, hematocrit or hemoglobin between WT and IFNγ-KO groups (**Supplementary Fig. S2C**).

Assessment of hepatic function revealed MHV infection significantly reduced total proteins (p < 0.0001), albumin (p < 0.0001 for WT and p < 0.01 for IFNγ-KO) and globulin (p < 0.0001) for both genotypes (**Fig. 1Ci**). These reductions were significantly less pronounced in the absence of IFNγ (p < 0.05 in the three cases) (**Fig. 1Cii**), as shown by post/pre ratios, indicative of reduced hepatic damage. No significant differences in total bilirubin, ALT, AST and GGT parameters were observed between genotypes post-infection (not shown). Regarding renal function, although no changes were observed in the BUN of WT animals, this parameter was significantly increased in the IFNγ-KO context (p < 0.0001) (**Fig. 1C**), suggesting affected kidney function. No changes were observed in the CRE levels of either group with the infection (**Fig. 1C**).

Concerning immune parameters, the infection caused a decrease in WBC total count and in lymphocytes percentage for both genotypes (p < 0.0001), while an increase in neutrophils, basophils, monocytes and eosinophils was observed in both cases after infection (p < 0.0001) (**Fig. 2Ai**). When comparing post/pre ratios between genotypes, IFNγ-KO mice presented a smaller reduction in lymphocytes (p < 0.001), less augmented basophils (p < 0.05) but a higher increase in neutrophils (**Fig. 2Aii**) (p < 0.05), showcasing a different immune strategy deployed in the absence of IFNγ. Regarding platelets and leukocyte ratios, both PLR and NLR changes were decreased in IFNγ-KO mice (p < 0.01) (**Fig. 2B**).

**Fig. 2.**
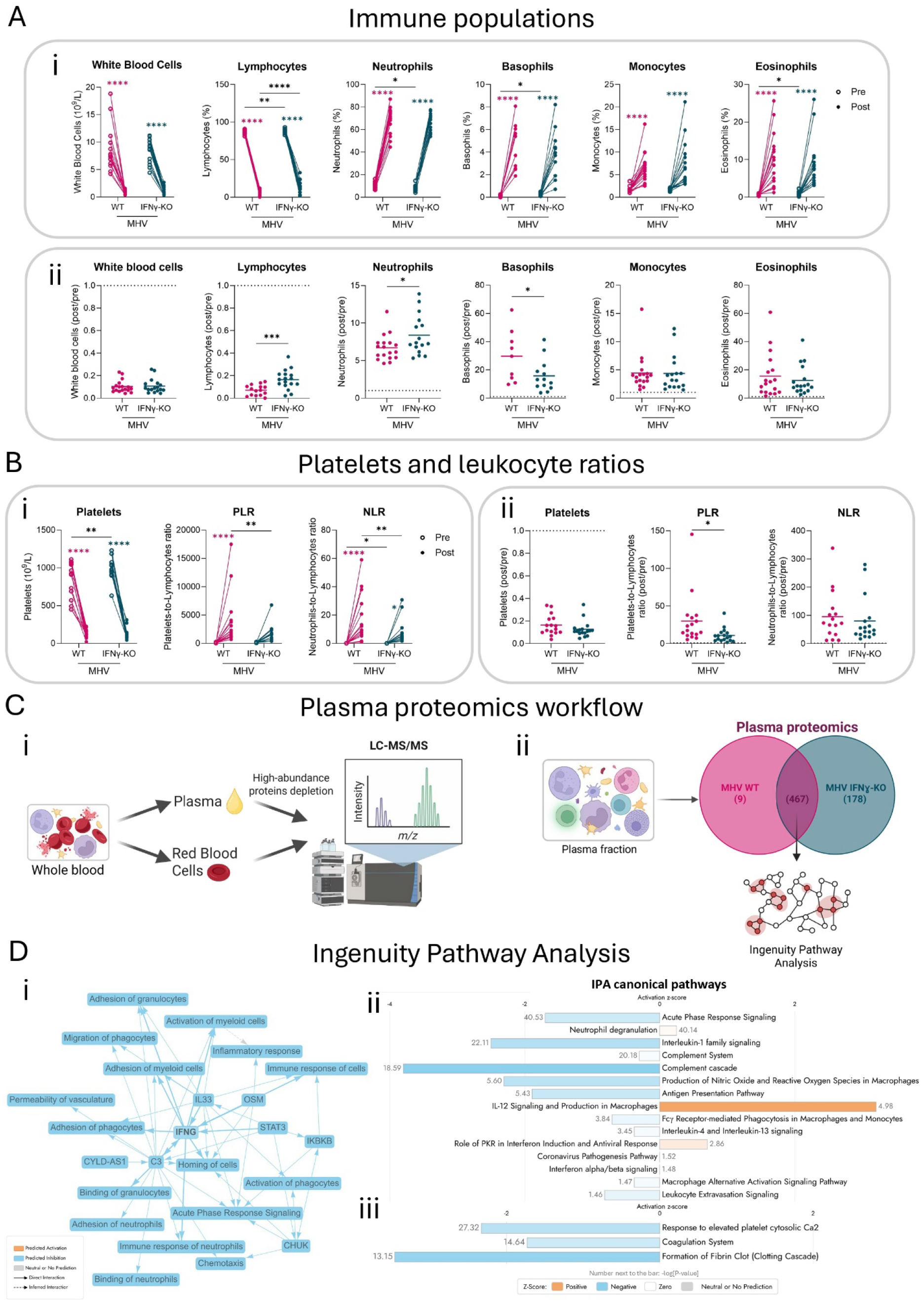
Plasma profiling following MHV infection. **(A)** Dot plots depicting immune cell population analysis following infection: total white blood cells (10^9^/L) and percentages (%) of each immune cell population (lymphocytes, neutrophils, basophils, monocytes and eosinophils), pre and post infection **(i)** and post/pre ratios **(ii)**. A post/pre ratio of 1 indicates no changes with the infection; ratios higher than 1 represent increases after infection; ratios lower than 1 represent decreases after infection. Comparing ratios between groups allows assessment of changes in the magnitude of the infection-induced response. **(B)** Dot plots depicting platelet counts (10^9^/L) and leukocyte ratios (platelets-to-lymphocytes and neutrophils-to-lymphocytes) pre and post infection **(i)** and post/pre ratios **(ii)**. Statistical significance was determined by mixed-effects analysis followed by Fisher’s LSD **(i)** or by Student’s t test **(ii)**, and was set at p < 0.05, with *p < 0.05; **p < 0.01; ***p < 0.001; ****p < 0.0001. **(C) (i)** Schematic representation of the proteomics experimental design. Briefly, blood was collected post-infection, plasma and red blood cells were fractioned and subjected to proteomics assessment by LC-MS/MS. Plasma fractions were depleted of albumin, transferrin and G immunoglobulins. Created with BioRender.com. **(ii)** Venn diagram of the number of proteins identified in the plasma fraction of WT and IFNγ-KO MHV-infected mice. The proteins shared between genotypes were subjected to differential expression and Ingenuity Pathway Analysis (IPA). Created with BioRender.com. **(D) (i)** IPA graphical summary plot of top predicted dysregulations observed in MHV-infected IFNγ-KO vs. MHV-infected WT plasma fractions. Orange = predicted activation; light blue = predicted inhibition; gray = neutral or no prediction; solid arrows = direct interaction; dashed arrows = inferred interaction. **(ii-iii)** Bar plots depicting the activation z-scores of IPA canonical pathways differentially activated in MHV-infected IFNγ-KO vs. MHV-infected WT plasma fractions. Significantly dysregulated pathways associated with inflammation **(i)** and coagulation **(ii)** are shown. Orange bars = positive z-scores; light blue = negative z-scores; white = zero z-scores; gray = neutral or no prediction; grey numbers next to bars represent −log(p-value). Statistical significance was set at -log[P-value] > 1.3.

Together, these findings indicate that IFNγ deficiency shapes a distinct clinical and immunological response to MHV infection, suggesting a less inflammatory state and decreased hypercoagulability.

### Proteomic profiling reveals enhanced systemic damage in the absence of IFNγ

We further explored infection-induced proteomics changes in blood. Whole blood was collected from MHV-infected WT and IFNγ-KO mice, and plasma and RBC were separated by centrifugation. Plasma was depleted of albumin, transferrin and G immunoglobulins, and proteins were purified from both compartments for proteomics analysis by LC-MS/MS (**Fig. 2Ci**).

A total of 654 proteins were confidently identified in plasma, with 467 of them shared between WT and IFNγ-KO groups, 178 unique to IFNγ-KO, and only 9 unique to WT (**Fig. 2Cii**). We performed differential expression analysis of the shared proteins (**Supplementary Table S1**), and the results were analyzed using Ingenuity Pathway Analysis (IPA) software. IFNγ deficiency was validated by this approach, and the graphical summary network (**Fig. 2Di)**, together with a canonical pathways analysis (**Fig. 2Dii**) predicted multiple inflammation-related categories to be inhibited in our IFNγ-KO model compared to WT. Moreover, the latter analysis also confirmed a negative regulation of coagulation and clotting in IFNγ-KO plasma (**Fig. 2Diii**). Plasma proteomics also predicted decreased infection and blood related diseases, such as anemia, blood clots, thrombus and proteinuria (**Supplementary Fig. S3**).

Overall, the plasma proteome revealed a markedly altered circulating landscape in the absence of IFNγ, highlighting reduced inflammatory and coagulation programs.

### IFNγ deficiency impairs RBC-mediated viral dissemination and antiviral responses

Given our previous observations of presence of viral particles in the blood (11), we sought to determine if IFNγ status affected viral dissemination mediated by RBC (**Fig. 3A**). Although no significant differences were observed in the viral load (**Fig. 3Bi**), the infectivity was reduced in the RBC fraction of IFNγ deficient mice (**Fig. 3Bii**).

**Fig. 3.**
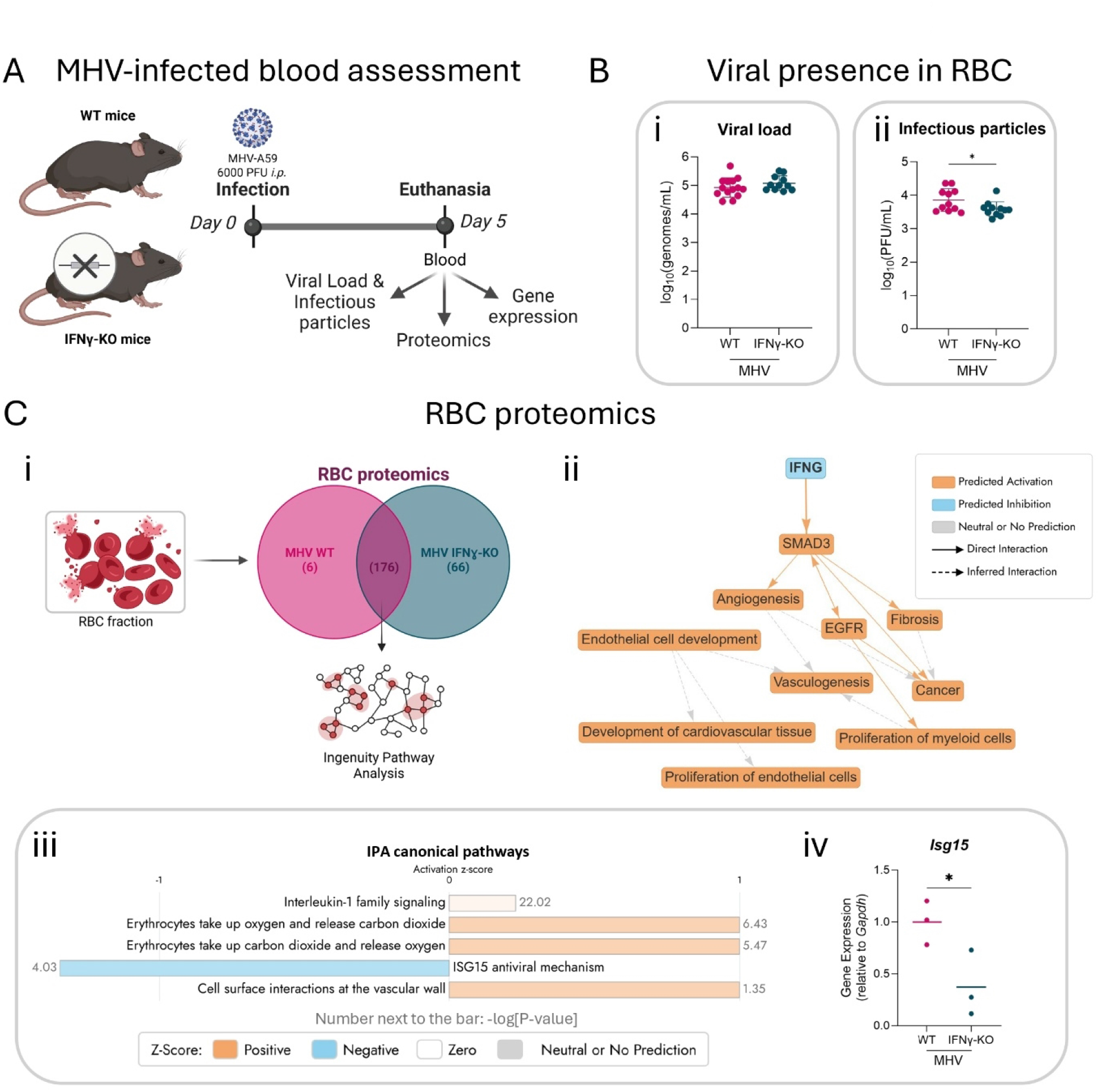
Systemic viral detection and erythrocyte proteomic alterations. **(A)** Schematic representation of the experimental design. Briefly, C57BL/6J wild-type (WT) and IFNγ-KO mice were infected i.p. with murine hepatitis virus (MHV) or mock (PBS). After 5 days, mice were euthanized and necropsy was performed. Blood was collected at baseline (pre) and post-infection (post) and samples were subjected to viral load and infectious particles determination, proteomics profiling and gene expression analyses. Created with BioRender.com. **(B)** Dot plots of viral presence in blood following infection. **(i)** Viral load quantification (log_10_ genomes/mL) was assessed by probed-qPCR and **(ii)** viral particles were assessed by plaque assays (log_10_(PFU/mL)) in the RBC fraction. **(C)** Red blood cells (RBC) fraction proteomic profiling. **(i)** Venn diagram of the number of proteins identified in the RBC fraction of WT and IFNγ-KO MHV-infected mice. The proteins shared between genotypes were subjected to differential expression and Ingenuity Pathway Analysis. Created with BioRender.com. **(ii-iii)** Ingenuity Pathway Analysis (IPA) of proteomics data. **(ii)** IPA graphical summary plot of top predicted dysregulations observed in MHV-infected IFNγ-KO vs. MHV-infected WT RBC fractions. Orange = predicted activation; light blue = predicted inhibition; gray = neutral or no prediction; solid arrows = direct interaction; dashed arrows = inferred interaction. **(iii)** Bar plot depicting the activation z-scores of IPA canonical pathways differentially activated in MHV-infected IFNγ-KO vs. MHV-infected WT RBC fractions. Significantly dysregulated pathways associated with RBC function are shown. Orange bars = positive z-scores; light blue = negative z-scores; white = zero z-scores; gray = neutral or no prediction; grey numbers next to bars represent −log(p-value). Statistical significance was set at -log[P-value] > 1.3. **(iv)** Isg15 expression levels in RBC fraction quantified by RT-qPCR, normalized to Gapdh, and presented as fold change compared to WT genotype. Statistical significance was calculated by Student’s t test and set at p < 0.05, with *p < 0.05.

To further investigate this phenomenon, we performed an RBC proteomics analysis to evaluate changes in erythrocyte-associated pathways and their potential contribution to viral dissemination. We identified a total of 248 proteins: 176 of them shared between genotypes, 66 exclusively observed in MHV IFNγ-KO mice and 6 solely in the WT genotype (**Fig. 3Ci** and **Supplementary Table S2**). The IPA summary confirmed IFNγ deficiency by predictions made from the differential protein expression, in association with increased SMAD3—a protein involved in brain injury protection (21)— fibrosis and angiogenesis (**Fig. 3Cii**). Among the pathways with predicted activation, IPA pointed to an enhanced RBC function and interactions at the vascular wall in MHV IFNγ-KO mice, while *Isg15* antiviral mechanism appeared to be repressed (**Fig. 3Ciii**). To confirm this prediction, we assessed *Isg15* expression by RT-qPCR and validated this impaired mechanism (**Fig. 3Civ**).

Collectively, the reduced RBC infectivity and the proteomic rewiring observed in IFNγ-deficient mice highlight a disrupted antiviral program.

### IFNγ deficiency promotes systemic viral dissemination but protects the brain

We next quantified viral burden across multiple organs using RT-qPCR and TCID_50_ assays (**Fig. 4Ai**). IFNγ-KO mice exhibited significantly increased viral load in tissues, including liver (p < 0.05), spleen (p < 0.05), heart (p < 0.05), and muscle (p < 0.05), compared to WT controls (**Fig. 4Aii**). Surprisingly, we found a significantly lower viral load in brain samples from IFNγ-KO mice, compared to WT controls (p < 0.01) (**Fig. 4Aii**). Consistently, infectious viral particle quantification confirmed the same trend observed with the viral RNA quantification (**Fig. 4Aiii-iv**).

**Fig. 4.**
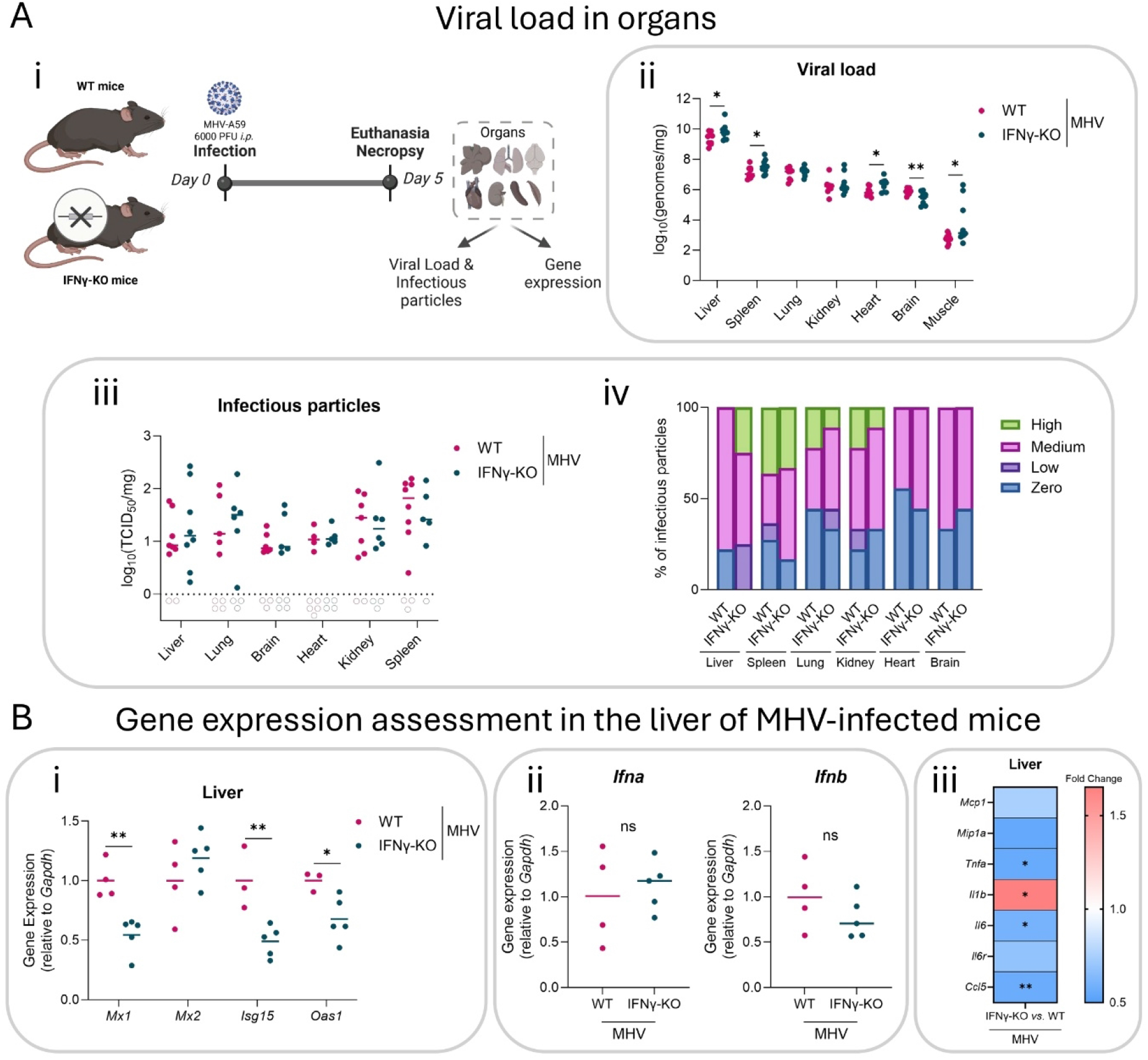
Effect of IFNγ deficiency in viral dissemination. **(A) (i)** Schematic representation of the experimental design. Briefly, C57BL/6J wild-type (WT) and IFNγ-KO mice were infected i.p. with murine hepatitis virus (MHV) or mock (PBS). After 5 days, mice were euthanized and necropsy was performed. Liver, spleen, lung, kidney, heart, brain and muscle were subjected to viral load and infectious particles determination, and gene expression analyses. Created with BioRender.com. **(ii)** Dot plot depicting viral load quantification (log_10_ genomes/mg) by probed-qPCR of liver, spleen, lung, kidney, heart, brain, and muscle from WT and IFNγ-KO MHV-infected mice. **(iii-iv)** Dot plot **(iii)** and stacked bar plot **(iv)** of infectious particles assessment by TCID_50_. Results (TCID_50_/mg) were categorized in zero, low, medium or high infectious particles by tertiles and % of mice in each category is shown **(iv)**. **(B)** Relative gene expression analysis in the liver of MHV-infected IFNγ-KO mice compared to MHV-infected WT controls. Dot plots **(i-ii)** and heatmap **(iii)** showing the expression levels of interferon-stimulated genes (ISGs) Mx1, Mx2, Isg15 and Oas1 **(i)**, type I interferons **(ii)** and inflammatory genes **(iii)** quantified by RT-qPCR, normalized to Gapdh, and presented as fold change compared to WT genotype. Statistical significance was calculated by Student’s t test and set at p < 0.05, with *p < 0.05; **p < 0.01.

Together, these findings demonstrate that IFNγ is required to restrict systemic viral replication in peripheral organs, while unexpectedly contributing to viral susceptibility in the brain.

### Antiviral and inflammatory gene expression is altered in IFNγ-KO Mice

We have previously shown the induction of the ISGs *MX1*, *MX2*, *ISG15* and *OAS1* in diverse models of Poly I:C treatment, SARS-CoV-2 and MHV-A59 infection (9). Thus, we explored the expression of these genes in our IFNγ-KO model of MHV infection. In the liver, the primary target organ of MHV, we observed a significant reduction in the expression of these ISGs: *Mx1* (p < 0.01), *Isg15* (p < 0.01) and *Oas1* (p < 0.05) (**Fig. 4Bi**). This impairment in the induction of these genes was attributed to the lack of IFNγ-KO, as there were no differences in the expression of *Ifna* or *Ifnb* between genotypes (**Fig. 4Bii**). Moreover, we also explored the expression of multiple cytokines, finding a significant reduction in the expression of the proinflammatory cytokines *Tnfa*, *Il6* and *Ccl5*, and increased levels of *Il1b* in the liver of infected IFNγ-KO mice (**Fig. 4Biii**).

Collectively, the impaired ISG induction and altered cytokine profile highlight a disrupted IFNγ-driven antiviral program in IFNγ-KO mice.

### IFNγ deficiency differentially affects viral burden across brain regions

Given the unexpected reduction of viral burden in the brain, we specifically examined proteomic signatures associated with neurological processes in our plasma proteomics results. Among the disease pathways with significant negative enrichment in plasma proteomics, we found categories related to neurological conditions and neuroinflammation, such as neuromuscular disease, brain lesions, neuronal cell death and activation of neuroglia (**Fig. 5A**).

**Fig. 5.**
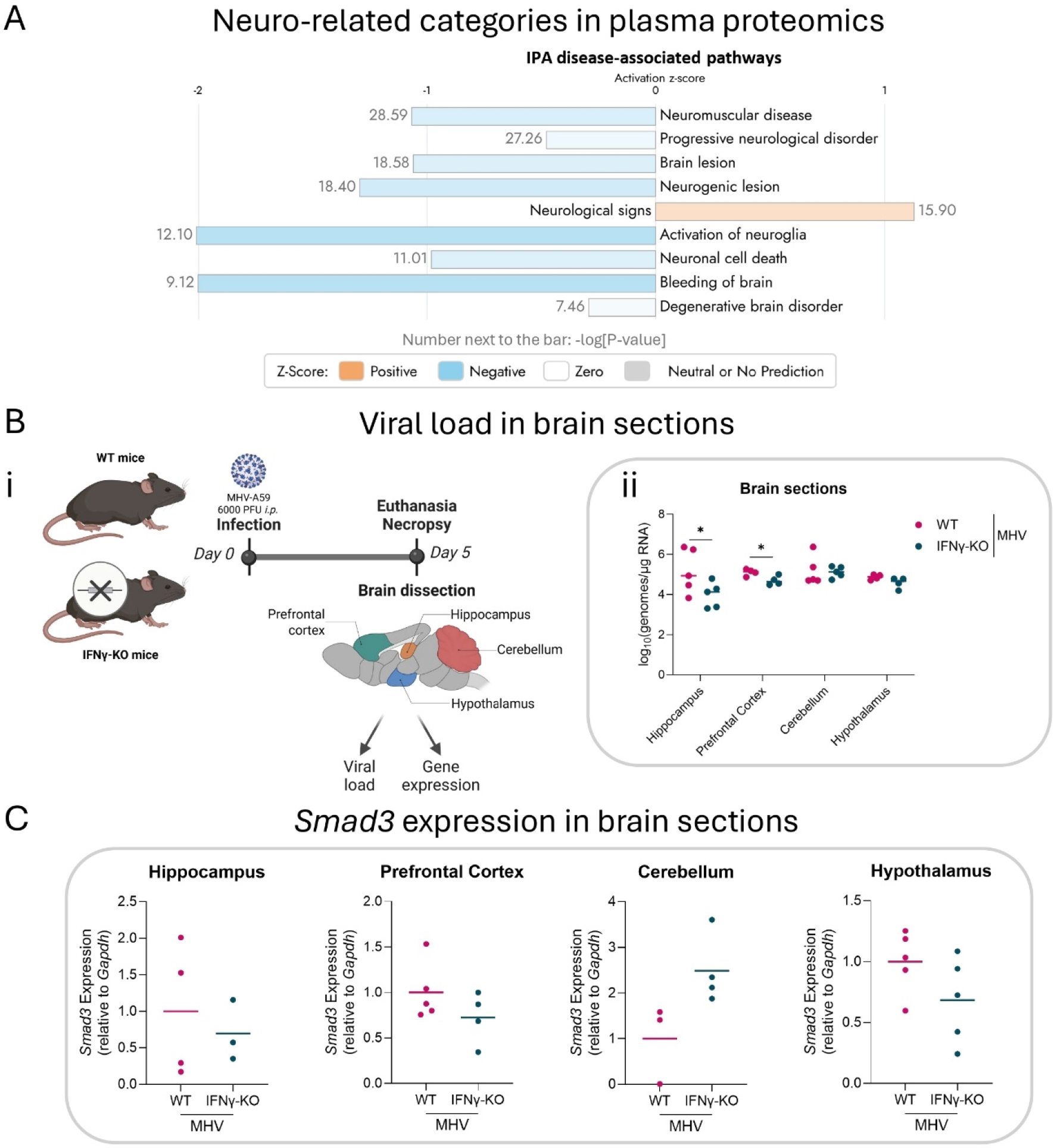
Neuroinfection and brain-region-specific susceptibility to MHV infection. **(A)** Bar plot depicting the activation z-scores of IPA disease-associated pathways differentially activated in MHV-infected IFNγ-KO vs. MHV-infected WT plasma fractions. Significantly dysregulated pathways associated with neurological diseases are shown. Orange bars = positive z-scores; light blue = negative z-scores; white = zero z-scores; gray = neutral or no prediction; grey numbers next to bars represent −log(p-value). Statistical significance was set at -log[P-value] > 1.3. **(B) (i)** Schematic representation of the experimental design. Briefly, C57BL/6J wild-type (WT) and IFNγ-KO mice were infected i.p. with murine hepatitis virus (MHV) or mock (PBS). After 5 days, mice were euthanized and necropsy was performed. Brains were obtained and further dissected in prefrontal cortex, hippocampus, hypothalamus and cerebellum sections, which were subjected to viral load determination and gene expression analyses. Created with BioRender.com. **(ii)** Dot plot depicting viral load quantification (log_10_ genomes/µg RNA) by probed-qPCR of hippocampus, prefrontal cortex, cerebellum and hypothalamus from WT and IFNγ-KO MHV-infected mice. **(C)** Smad3 expression levels in hippocampus, prefrontal cortex, cerebellum and hypothalamus sections quantified by RT-qPCR, normalized to Gapdh, and presented as fold change compared to WT genotype. Statistical significance was calculated by Student’s t test and set at p < 0.05, with *p < 0.05.

To further explore these observations, we dissected the brains after infection and measured viral load by RT-qPCR in the prefrontal cortex, hippocampus, hypothalamus and cerebellum of IFNγ-KO and WT mice (**Fig. 5Bi**). Viral load was significantly decreased in the prefrontal cortex and hippocampus of IFNγ-KO mice (p < 0.05), while no differences were observed in hypothalamus or cerebellum (**Fig. 5Bii**). These data indicate a region-specific role of IFNγ in modulating viral tropism.

Since SMAD3, a protein involved in protection against cortical damage (21) and a mediator of endothelial permeability and neuroinflammation in viral and sterile inflammatory contexts (21–23), was predicted as activated in RBC proteomics (**Fig. 4Cii**), we assessed its expression across brain regions. We observed a trend toward reduced SMAD3 expression in the hippocampus, prefrontal cortex and hypothalamus of IFNγ-KO mice, whereas an opposite trend was detected in the cerebellum (**Fig. 5C**).

Altogether, the data suggest that the absence of IFNγ shapes a region-dependent neuroprotective landscape, consistent with the reduced viral load observed in cortical areas.

### IFNγ shapes neuronal antiviral and neuroinflammatory programs in human COVID-19 brain datasets

To determine whether IFNγ-related transcriptional programs were also engaged in the human brain during coronavirus infection, we analyzed publicly available transcriptomic datasets from neurons treated with IFNγ and frontal cortex of COVID-19 patients.

We examined the impact of IFNγ on neuronal transcriptional responses by analyzing RNA-seq data from primary cultures of neurons treated with IFNγ and we performed differential expression analysis and IPA (**Fig. 6Ai** and **Supplementary Table S3**). The graphical summary revealed that treatment of neuron cultures with IFNγ increased the expression of genes related to neuroinflammation and antiviral response (**Fig. 6Aii**). Among the disease-associated expression pattern, we detected similar dysregulations as the ones observed in different brain lesions and disorders (**Fig. 6Aiii)**, while neuroinflammation and, surprisingly, transcriptional activity of SMAD2/SMAD3:SMAD4 heterodimer appeared significantly dysregulated among the altered canonical pathways (**Fig. 6Aiv**). Moreover, when analyzing potential upstream regulators involved in IFNγ-induced differential profiles, SMAD3 was the second most activated of the SMAD family (**Fig. 6Av**). These findings further support a diminished neuroinflammatory environment in the absence of IFNγ and point out to a SMAD3-mediated mechanism.

**Fig. 6.**
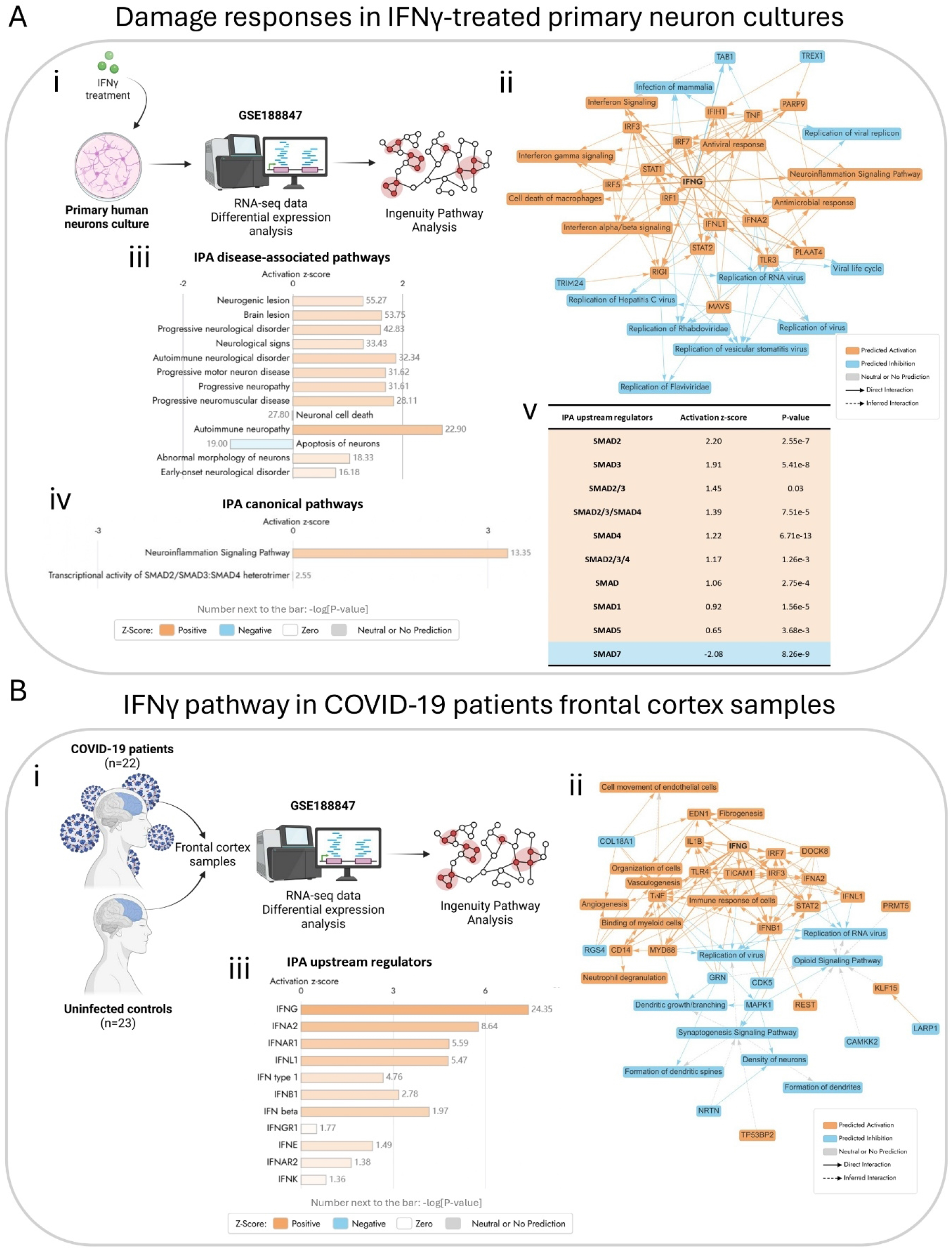
IFNγ pathway activation in human COVID-19 and neuronal damage modeling. **(A) (i)** Schematic representation of the bioinformatics analysis. Briefly, we used transcriptomics data of primary human neuron cultures treated with IFNγ and untreated controls available in GSE188847. We performed differential expression and Ingenuity Pathway Analysis (IPA). Created with BioRender.com. **(ii)** IPA graphical summary plot of top predicted dysregulations observed in IFNγ-treated vs. untreated primary human neuron cultures. Orange = predicted activation; light blue = predicted inhibition; gray = neutral or no prediction; solid arrows = direct interaction; dashed arrows = inferred interaction. **(iii-iv)** Bar plot depicting the activation z-scores of neurological disease **(iii)** and canonical neuroinflammation and SMAD transcriptional activity **(iv)** pathways in IFNγ-treated vs. untreated primary human neuron cultures. Orange bars = positive z-scores; light blue = negative z-scores; white = zero z-scores; gray = neutral or no prediction; grey numbers next to bars represent −log(p-value). **(v)** Table depicting SMAD molecules significantly identified as IPA upstream regulators. **(B) (i)** Schematic representation of the bioinformatics analysis. Briefly, we used transcriptomics data of frontal cortex samples from COVID-19 patients and uninfected controls available in GSE188847. We performed differential expression and Ingenuity Pathway Analysis (IPA). Created with BioRender.com. **(ii)** IPA graphical summary plot of top predicted dysregulations observed in COVID-19 vs. uninfected human frontal cortex samples. Orange = predicted activation; light blue = predicted inhibition; gray = neutral or no prediction; solid arrows = direct interaction; dashed arrows = inferred interaction. **(iii)** Bar plot depicting the activation z-scores of interferons as upstream regulators in COVID-19 vs. uninfected human frontal cortex samples. Orange bars = positive z-scores; light blue = negative z-scores; white = zero z-scores; gray = neutral or no prediction; grey numbers next to bars represent −log(p-value). Statistical significance was set at -log[P-value] > 1.3.

To bridge our findings with human disease, we analyzed a publicly available transcriptomic dataset from COVID-19 patients’ frontal cortexes (**Fig. 6B** and **Supplementary Table S4**). Differential expression and pathway analysis revealed predicted activation of the IFNγ signaling pathway in the frontal cortex samples of COVID-19 patients, in association with increased fibrogenesis and immune response, and decreased viral replication and formation of neural networks (**Fig. 6Bii**). Among the different interferons, IFNγ was the one with the highest activation z-score in the frontal cortex from COVID-19 patients (**Fig. 6Biii**), supporting that IFNγ–driven responses are active in the human brain during SARS-CoV-2 infection and may contribute to the observed neuropathology.

Taken together, the IFNγ-induced transcriptional programs in neurons and the strong activation of IFNγ-responsive pathways in COVID-19 frontal cortex converge to show that IFNγ is a key driver of neuronal antiviral and neuroinflammatory responses, including modulation of SMAD-dependent signaling.

## Discussion

Herein, using a preclinical mouse model, we reveal a dual and tissue specific role of IFNγ during MHV infection, simultaneously controlling systemic viral dissemination while promoting brain susceptibility and neuroinflammatory signatures. Using a combination of a preclinical IFNγ-KO infection model, proteomics, and bioinformatics analyses of public transcriptomics data, we show that IFNγ is a central determinant of coronavirus pathogenesis. IFNγ deficiency increased viral burden in peripheral organs but markedly reduced viral load in multiple brain regions, including the prefrontal cortex and hippocampus, uncovering a previously unrecognized role for IFNγ in facilitating coronavirus neuroinvasion.

Interferons, key regulators of antiviral immunity, orchestrate innate and adaptive responses to viral infections including coronaviruses (24). IFN-I constitute the canonical first line of defense, inducing hundreds of ISGs that restrict viral replication (25). However, coronaviruses (including SARS-CoV, MERS-CoV, SARS-CoV-2, and MHV) have evolved potent IFN-I evasion strategies (26–29), resulting in delayed or blunted IFN-I responses, a defining feature of severe coronavirus infections (30). Impaired IFN-I signaling is strongly associated with increased viral load, hyperinflammation, and mortality (27,31–33).

In contrast, IFNγ (type II IFN) has been widely disregarded in the context of coronavirus infections (34), exhibiting controversial roles in both immunomodulation and immunotolerance (35). Despite increased viral loads, our IFNγ-deficient infected mouse model exhibited attenuated systemic damage, including reduced alterations in hepatic function and decreased activation of coagulation pathways. Plasma proteomics revealed downregulation of inflammatory and thrombotic signatures in IFNγ-KO animals, consistent with reports linking interferon-driven inflammation to coagulopathy and multiorgan dysfunction (36). The decreased PLR and NLR observed in IFNγ-KO mice further supports less severe inflammatory and thrombotic state, as elevated PLR and NLR have been associated with poor outcomes in viral infections including COVID-19 (37,38). Further, our findings reinforce the concept that IFNγ deficiency resulted in reduced ISG induction and increased systemic viral dissemination despite preserved IFN-I expression, highlighting the non-redundant antiviral role of IFNγ. These results align with evidence from SARS-CoV-2 and other coronavirus infections showing that excessive cytokine production contributes to tissue damage and disease severity (39,40).

Beyond systemic pathology, neurological manifestations are increasingly recognized in acute COVID-19 and Long-COVID, including anosmia, cognitive impairment, encephalopathy, neuroinflammation, and neurovascular dysfunction (41,42). Severe disease is characterized by elevated circulating cytokines such as IL-6, TNF-α, and IFNγ, contributing to blood–brain barrier (BBB) disruption and neuroinflammatory cascades (43). Moreover, IFNγ has been noted as a driver of BBB permeability in association with several neurological infections (44). In our study, IFNγ–deficient mice exhibited significantly lower viral load in the prefrontal cortex and hippocampus, contrasting with the increased viral burden observed in peripheral organs.

The role of IFNγ in viral neuropathogenesis is not unique to coronaviruses and appears to be highly context dependent across neurotropic infections. In West Nile Virus (WNV) infection, IFNγ contributes to viral control within the CNS, and its deficiency is associated with increased viral burden and susceptibility to neuroinvasive disease, supporting a predominantly protective role (45). Conversely, elevated IFNγ expression has been reported in severe neurological manifestations of Zika virus infection, including microcephalic fetal brains, where heightened IFNγ signaling has been linked to inflammatory responses and neurodevelopmental abnormalities (46). Similar dual functions have been described in other viral and bacterial infections of the CNS, such as pneumococcal meningitis, where IFNγ is required for efficient clearance but may simultaneously promote neuroinflammation, neuronal dysfunction, BBB alterations, and long-term neurological sequelae (47–49). Together, these observations suggest that the biological consequences of IFNγ signaling in the brain depend on the balance between protection and immune-mediated tissue injury. Our findings extend this paradigm to coronavirus infection, identifying IFNγ as a factor that may simultaneously restrict systemic viral dissemination while enhancing neurovascular and neuroinflammatory processes that favor brain involvement.

An additional layer of complexity tackled in our work encompasses the possible routes of entry of coronavirus to the central nervous system. It has previously been proposed that neurovirulence of coronavirus could be explained by nerve infection and hematogenous invasion (42). In the latter, points of weakness in the BBB may arise from viral-induced release of inflammatory mediators and cytokine storms leading to destabilization or disruption of BBB tight-junctions (50). Moreover, the “trojan horse” mechanism has been described in leukocytes and myeloid cells (50). Our previous work has shown MHV presence in the RBC of infected mice, suggesting a mechanism of “hitchhiking” dissemination that could bypass classical immune surveillance (11). Here, although viral RNA levels in RBC fractions were comparable between genotypes, infectious viral particles were significantly reduced in IFNγ–deficient mice. These results indicate that IFNγ may play a role in these RBC-associated particles, potentially explaining the differences in brain viral load.

Consistent with these observations, RBC proteomics further revealed IFNγ–dependent regulation of pathways related to endothelial interaction, vasculogenesis and cell surface processes, suggesting that IFNγ may enhance the capacity of RBCs to interact with vascular endothelium and potentially deliver viral particles to susceptible tissues. Notably, *ISG15* was downregulated in IFNγ-deficient RBCs, providing a validation of our proteomic findings. Although ISG15 is classically considered a type I interferon-stimulated gene (51), its reduced expression in IFNγ-KO mice despite preserved IFN-I expression suggests a more complex regulatory network in which IFNγ contributes directly or indirectly to ISG15 induction. This observation reinforces the concept that IFNγ signaling cannot be viewed merely as a parallel antiviral pathway, but rather as an important modulator of canonical interferon-responsive programs. The regulation of *ISG15* by IFNγ may therefore represent an additional mechanism linking IFNγ activity to coronavirus dissemination and tissue tropism.

In COVID-19, BBB breakdown has been documented in both patients and animal models, correlating with microglial activation, neuroinflammation, and viral RNA detection in brain tissue (43). Additionally, IFNγ was reported to generate BBB leakage during reovirus infection through Rho kinase-mediated mechanisms (52). Our findings that IFNγ deficiency reduces brain viral load despite increased systemic dissemination suggest that IFNγ may facilitate viral access to the brain by simultaneously enhancing RBC-mediated vascular interactions and weakening BBB integrity. This interpretation is further supported by the predicted activation of IFNγ, neurovascular and neuroinflammatory pathways, and neuronal injury signatures in IFNγ–treated neurons and COVID-19 frontal cortex samples, as well as the upregulation of SMAD3. The brain-specific protection in our *in vivo* IFNγ-KO model was accompanied by decreased activation of neuro-related pathways in IFNγ–deficient plasma proteomics and by reduced *SMAD3* expression, supporting SMAD3 as a mechanistic bridge between IFNγ signaling and coronavirus-induced brain pathology.

Collectively, our results demonstrate that IFNγ plays a dual role during coronavirus infection. It is essential for controlling systemic viral replication and limiting multiorgan damage, yet it simultaneously contributes to viral susceptibility and inflammatory processes in the brain, possibly mediated by SMAD3-dependent modulation of neurovascular and neuroimmune pathways. These findings underscore the need for carefully balanced therapeutic strategies targeting interferon pathways, where modulation or partial inhibition may be required to optimize antiviral efficacy while minimizing host mediated pathology. Based on our findings, neurologically focused attenuation of IFNγ signaling (through JAK1/2 inhibition, anti-IFNγ antibodies or localized delivery of IFNγ antagonists) may represent a promising approach to prevent or mitigate coronavirus associated neurological complications, while preserving the systemic antiviral benefits of IFNγ.

### Limitations of the study

While our data establish strong associations between IFNγ signaling and tissue-specific outcomes, causal mechanisms particularly in the brain, require further investigation. Particularly, the IFNγ-SMAD3 axis should be further characterized.

IPA is a powerful application that enables researchers to integrate their studies with an expert-curated knowledge base, including causal relationships identified across publications. However, pathway predictions derived from IPA should be validated experimentally in this specific experimental setting. Moreover, although the MHV-A59 model recapitulates key aspects of coronavirus biology, differences with others virus from the same family must be considered.

Brain dissections are difficult to perform and RNA yield is not always sufficient. Thus, our samples were limited and bigger experiments should be carried out to continue assessing gene expression and find statistically significant differences.

Future studies should explore spatiotemporally controlled IFNγ interventions to fine-tune the balance between antiviral immunity and neuroprotection and determine its clinical utility.

## Acknowledgments

All schematic representations were created using BioRender.com. The summary networks, upstream regulators/canonical pathways/disease pathways plots were generated through the use of QIAGEN IPA (QIAGEN Inc., https://digitalinsights.qiagen.com/IPA) and QIAGEN IPA Interpret (QIAGEN Inc., https://digitalinsights.qiagen.com/products-overview/discovery-insights-portfolio/analysis-and-visualization/qiagen-ipa/qiagen-ipa-interpret/).

## Authorship contributions

**Conceptualization:** AS, JMC, GM, MC, GG, AT.

**Methodology:** AS, APA, PP, PS, GP, JLP, RS, MP, MPV, JMC, PM, GM, MC, GG, AT.

**Investigation:** AS, JC, EV, JMC, GM, MC, GG, AT.

**Visualization:** AS, PS, JMC, GM, MC, GG, AT.

**Statistical Analyses:** AS, PS, GP, RS, JC, GG, AT.

**Supervision:** MPV, JMC, PM, GM, MC, GG, AT.

**Writing—original draft:** AS, GG, AT.

**Writing—review & editing:** AS, APA, PP, PS, GP, EV, EL, JMC, GM, MC, GG, AT.

**Project administration:** PM, GM, MC, GG, AT.

**Funding acquisition:** AS, EV, EL, PM, GM, MC, GG, AT.

All authors agreed to submit the manuscript and have read and approved its final version. Authors take full responsibility for its content.

## Supporting information captions

### Supplementary Fig.s

**Supplementary Fig. S1. Baseline characterization of IFNγ knockout mice. (A)** Schematic representation of the experimental design for baseline characterization of IFNγ knockout (IFNγ-KO) and C57BL/6J wild-type (WT) mice under uninfected conditions. Organs and blood were collected. Biochemical and hematological parameters were analyzed, and organs were weighted. Created with BioRender.com. **(B)** Dot plot depicting the body weight (g) comparison between WT and IFNγ-KO mice. **(C)** Dot plots depicting the comparison of organ weights between WT and IFNγ-KO mice. **(D)** Dot plots depicting biochemical parameters including hepatic (total proteins, albumin, globulin, aspartate aminotransferase, alanine aminotransferase and gamma-glutamyl transpeptidase) **(i)** and renal (blood urea nitrogen and creatinine) **(ii)** function markers, measured in WT and IFNγ-KO mice. **(E) (i)** Dot plot depicting the flow cytometry analysis of total white blood cells in WT and IFNγ-KO mice. **(ii)** Bar plot showing the percentage of each immune cell population (lymphocytes, neutrophils, basophils, monocytes and eosinophils), measured in WT and IFNγ-KO mice. **(F)** Dot plots depicting the hematological parameters measured in WT and IFNγ-KO mice, including red blood cells number, hematocrit, hemoglobin, platelets and leukocyte ratios (platelets-to-lymphocytes and neutrophils-to-lymphocytes). Data are presented as mean and dispersion. Statistical analysis was performed using Student’s t test and was set at p < 0.05, with *p < 0.05; **p < 0.01.

**Supplementary Fig. S2. Physiological and hematological alterations upon MHV infection. (A)** Schematic representation of the experimental design. Briefly, C57BL/6J wild-type (WT) and IFNγ-KO mice were infected *i.p.* with murine hepatitis virus (MHV) or mock (PBS). After 5 days, mice were euthanized and necropsy was performed. Blood was collected at baseline (pre) and post-infection (post). Biochemical and hematological parameters were analyzed, and organs were weighted. Created with BioRender.com. **(B)** Dot plots comparing organ weights (% of body weight) of MHV-infected and mock-infected WT and IFNγ-KO mice. **(C)** Dot plots of the blood parameters, including total red blood cells (10^12^/L), hematocrit (%) and hemoglobin (g/L) post/pre ratios in WT and IFNγ-KO MHV-infected mice. A post/pre ratio of 1 indicates no changes with the infection; ratios higher than 1 represent increases after infection; ratios lower than 1 represent decreases after infection. Comparing ratios between groups allows assessment of changes in the magnitude of the infection-induced response. Statistical significance was determined by **(B)** one-way ANOVA followed by post-hoc Tukey’s test, **(C)** mixed-effects analysis followed by Fisher’s LSD **(i)** or by Student’s t test **(ii)**, and was set at p < 0.05, with *p < 0.05; **p < 0.01; ***p < 0.001; ****p < 0.0001.

**Supplementary Fig. S3. Confirmation of decreased infection and coagulation in plasma proteomics. (A)** Bar plots depicting the −log(p-value) of Ingenuity Pathway Analysis (IPA) disease-associated pathways differentially activated in MHV-infected IFNγ-KO vs. MHV-infected WT plasma fractions. Significantly dysregulated pathways associated with viral infections **(A)** and blood diseases **(B)** are shown. Orange bars = positive z-scores; light blue = negative z-scores; white = zero z-scores; gray = neutral or no prediction. Statistical significance was set at p < 0.05.

### Supplementary Tables

**Supplementary Table S1. Differential abundance of plasma proteomics.**

**Supplementary Table S2. Differential abundance of RBC proteomics.**

**Supplementary Table S3. Differential expression of IFNγ-treated primary human neuron cultures.**

**Supplementary Table S4. Differential expression of COVID-19 patients’ frontal cortexes.**

